# A midnight snack for spirulina: a dual-substrate kinetic model for second cheese whey supplementation of *Arthrospira platensis*

**DOI:** 10.64898/2026.11.19.767785

**Authors:** Claudio Brandoli, Antonio Spanu, Giulia Neri, Francesca Masino, Elisabetta Sgarbi, Leonardo Miele

## Abstract

Arthrospira platensis (Spirulina) is a widely used strategic cyanobacterium whose cultivation cost remains a primary barrier to large-scale production. Mixotrophic growth on dairy by-products offers a simultaneous route to increase biomass concentration and valorize industrial waste streams. Here, we investigate the effect of Parmigiano Reggiano second cheese whey supplementation on *A. platensis* batch cultivation under continuous and alternating photoperiods. Whey supplementation increased final biomass in a dose-dependent manner under both continuous and alternating light. At the higher supplementation level, cultures under alternating light reached the same final biomass as under continuous illumination within the cultivation window. To interpret the interaction between whey availability, photoperiod, and biomass accumulation, we developed a kinetic model in which inorganic substrate assimilation is light-dependent, while whey-associated growth can contribute during both light and dark phases through a phenomenological dark-phase coefficient. Model and experimental results are consistent with the assumption that photoperiod mainly modulates the rate of biomass accumulation rather than the attainable biomass, under sufficient substrate availability. These results support a nutrient-stoichiometric interpretation of the biomass ceiling and suggest that second cheese whey supplementation can reduce dependence on continuous artificial lighting, providing a promising route for more resource-efficient *Arthrospira* cultivation.

**Highlights:**

- Second cheese whey increased *Arthrospira platensis* biomass up to 1.8-fold.
- At 2% whey, alternating-light matched continuous-light biomass production.
- A dual-substrate model separates light-driven and whey-driven growth.

## 1. Introduction

Cyanobacteria strains of the species *Arthrospira platensis* Gomont [1], commonly known as Spirulina, have gained renewed attention in recent decades due to their biotechnological potential [2], rapid growth rate [3], and high photosynthetic efficiency [4], which make them attractive for large-scale biotechnological applications [5]. Adapted to highly alkaline and hypersaline environments [6], *A. platensis* is characterized by a biochemical composition rich in proteins and other high-value compounds, supporting its widespread use in the food, feed, and nutraceutical sectors [7, 8]. Under optimal cultivation conditions, protein content can exceed 60% of dry biomass, positioning Spirulina among the most protein-rich microbial resources available and as a promising sustainable alternative to conventional animal-derived protein sources [9, 10, 11].

Beyond its nutritional value, Spirulina is increasingly recognized as a versatile biomass for sustainable agricultural and industrial applications [12, 13]. Its biomass productivity can substantially exceed that of conventional terrestrial crops [14], while requiring considerably less land and freshwater than livestock-based protein production [15]. These characteristics have also promoted its use in a broad range of fields, including agricultural biostimulants, biofuels, pharmaceuticals, cosmetics, and environmental remediation [16, 17, 18, 19].

A persistent barrier to scaling production is its relatively high cultivation cost. Furthermore, conventional autotrophic open-air systems remain constrained by limited nutrient availability, non-uniform light distribution and environmental fluctuations, leading to variable biomass yield and biochemical composition [20, 21]. In closed and artificially illuminated systems in particular, lighting is a dominant contributor to operating cost and energy demand [22], so any strategy that relaxes the dependence of productivity on continuous illumination — for example by exploiting natural light–dark cycles or reduced illumination schedules — has direct economic and sustainability implications. Mixotrophic cultivation, which combines photosynthetic carbon fixation with the assimilation of exogenous organic substrates, can improve biomass yield while enabling the valorization of nutrient-rich industrial effluents, thereby supporting circular bioeconomy strategies [5, 23, 24, 25]. In particular, the dairy industry annually generates large volumes of whey-derived by-products, whose high organic load and residual nutrient content make their management economically and environmentally challenging [26, 27].

Whey-derived dairy by-products have emerged as promising substrates for mixotrophic cyanobacterial cultivation [28, 29, 30]. Among these, second cheese whey remains comparatively underexplored despite retaining lactose, minerals, and other potentially valuable nutrients [31]. In some areas of Northern Italy, where Parmigiano Reggiano cheese production is highly concentrated [32], substantial amounts of second cheese whey are generated as a by-product of the ricotta manufacturing process, making its valorization particularly relevant within regional circular bioeconomy frameworks [33].

Despite the growing body of literature on mixotrophic cultivation of *A. platensis*, several aspects related to process optimization and predictive modelling remain insufficiently explored [34, 35, 36]. Most experimental studies have evaluated the effect of selected cultivation conditions on final biomass concentration and biochemical composition, while dynamic interactions between culture substrate availability, light regime, and growth kinetics remain less frequently investigated [28, 29, 30]. First, the dose-dependent effect of dairy supplementation on the maximum attainable biomass remains insufficiently characterized, as most studies evaluate single substrate concentrations and report endpoint biomass values with limited integration of substrate conversion stoichiometry. Second, kinetic models applied to *A. platensis* mixotrophic growth have often relied on simplified single-substrate Monod-type formulations [34, 37], where growth limitation is typically described as a function of a single substrate concentration. Such approaches may not fully capture the coupled effects of organic carbon availability and light regime, which jointly regulate photosynthetic activity, metabolic fluxes, and biomass production.

To address these knowledge gaps, this study investigated the cultivation of *A. platensis* on Parmigiano Reggiano second cheese whey, a dairy by-product generated in large quantities in Northern Italy, at two whey supplementation levels and under two photoperiod regimes. The resulting biomass growth dynamics were described using a dual-substrate, light-dependent kinetic model calibrated through Bayesian Markov chain Monte Carlo (MCMC) inference. Two central questions were addressed: (i) does whey supplementation increase the maximum attainable biomass concentration in a dose-dependent and quantitatively predictable manner? (ii) Can organic substrate availability reduce the dependence of biomass accumulation on photoperiod without compromising final biomass concentration? Answering these questions may contribute to the development of more sustainable and lighting-efficient strategies for industrial-scale *Arthrospira* cultivation.

## 2. Materials and Methods

### 2.1. Culture conditions

The *A. platensis* strain RCC6351 used in this research was obtained from the Roscoff Culture Collection (RCC, France; https://roscoff-culture-collection.org/) via the Unione Spirulina Biologica Italiana Association (Italy; https://unionespirulina.it/). Cyanobacterium strain was cultivated in BG11 medium (Blue Green Medium, PhytoTech Labs; https://phytotechlab.com/), as described by Marrez et al. [38], following standard techniques for *in vitro* plant cell autotrophic cultures. The initial pH of the cultures was adjusted to 8.0, a condition reported to promote optimal biomass concentration and physiological performance in *A. platensis* [39, 40]. Medium was autoclaved at 121°*C*, 1 atm, for 20 minutes before inoculation and flasks were always maintained covered with cotton plugs to prevent contamination. The detailed composition of the BG11 medium is reported in Table 1.

**Table 1.** Composition of BG11 culture medium.

| **Component** | **Concentration (g L<sup>−1</sup>)** |
|---|---|
| NaNO<sub>3</sub> | 1.5 |
| CaCl<sub>2</sub> · 2H<sub>2</sub>O | 0.036 |
| MgSO<sub>4</sub> · 7H<sub>2</sub>O | 0.075 |
| K<sub>2</sub>HPO<sub>4</sub> | 0.04 |
| Citric acid | 0.006 |
| (NH<sub>4</sub>)<sub>5</sub>[Fe(C<sub>6</sub>H<sub>4</sub>O<sub>7</sub>)<sub>2</sub>] | 0.006 |
| Na<sub>2</sub>CO<sub>3</sub> | 0.02 |
| Na<sub>2</sub>-EDTA | 0.001 |
| H<sub>3</sub>BO<sub>3</sub> | 0.0028 |
| ZnSO<sub>4</sub> · 7H<sub>2</sub>O | 0.00022 |
| MnCl<sub>2</sub> · 4H<sub>2</sub>O | 0.0018 |
| NaMoO<sub>4</sub> · 2H<sub>2</sub>O | 0.00039 |
| CuSO<sub>4</sub> · 5H<sub>2</sub>O | 0.00008 |
| Co(NO<sub>3</sub>)<sub>2</sub> · 6H<sub>2</sub>O | 0.00005 |

Cultivation was carried out under controlled environmental conditions inside a climate growth chamber (Frigor-Box, series 900/1200). The flasks were randomly distributed across the shelves of the growth chamber and systematically rotated between positions throughout the cultivation period to minimize potential positional effects associated with differences in light exposure and other local environmental conditions. A cultivation temperature of 25 ± 1°*C* was selected to reflect a temperature relevant to industrial-scale *A. platensis* cultivation, particularly in temperate regions, while limiting the need for active temperature control. Illumination was provided by white led lamps positioned 32 *cm* above the cultures. The tested conditions included:

- 25 ± 1° 69 *C*, 16 hours light/8 hours dark photoperiod
- 25 ± 1° 71 *C*, continuous light (24 hours)

Light intensity was measured with a photoradiometric probe (Delta OHM, HD2102.21) and set to 105 ± 2 *μ*mol photons *m*^−2^*s*^−1^, within the range previously investigated for *A. platensis* [41]. This controlled irradiance ensured uniform and consistent light exposure across samples.

Cultures were grown for one week in 250 mL Erlenmeyer flasks containing 100 *mL* of BG11 medium and inoculated with 10 *mL* of Spirulina at a concentration of 0.1 *g L*^−1^ before being used for the experiments. This pre-incubation period allowed the cells to reach a physiologically active state, reducing the lag phase and promoting uniform growth in subsequent experiments, as suggested by Pelizer et al. [42], who reported that inocula of 6-day-old

#### A. platensis colonies showed optimal growth performance

Mixotrophic conditions were obtained by adding 1% and 2% (v/v) of Parmigiano Reggiano second cheese whey (SCW henceforth), supplied by Nuova Latteria Fontana (Rubiera, Reggio Emilia, Italy), to BG11, which served as standard culture medium. Physicochemical characterization of SCW (Table 2) was performed before its use as a cultivation substrate. Moisture and dry matter were determined by gravimetric drying at 100°*C* until constant weight, whereas ash content was measured by incineration at 550°*C*. Lipids were extracted according to the method described by Folch et al. [43], protein content was determined using the Lowry method [44] and soluble solids were measured by refractometry and expressed as °Brix. Since lactose is the predominant soluble carbohydrate typically found in SCW, this value was considered an indirect indicator of the lactose-containing soluble fraction, although refractometry measures overall soluble solids rather than lactose specifically. Results are reported as mean ± standard deviation.

**Table 2.**
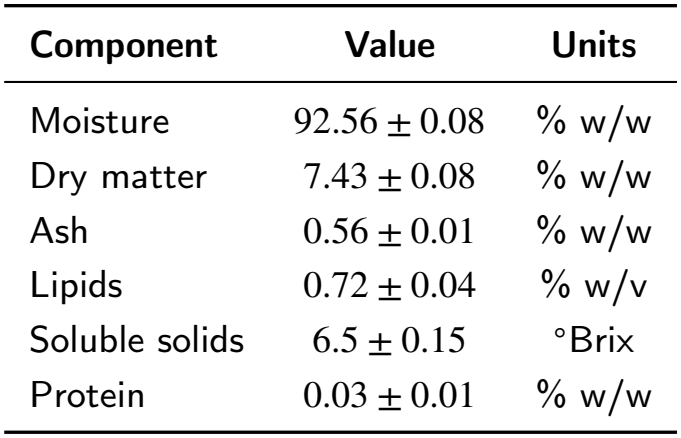
Physicochemical characteristics of Parmigiano Reggiano second cheese whey used as mixotrophic substrate.

### 2.2. Experimental design and growth measurements

Cultures were sampled every 5 days for 25 days, corresponding to six sampling points (T0, T5, T10, T15, T20, and T25). To minimize sampling bias due to the aggregative nature of Spirulina, mediated by extracellular polymeric substances excreted on the surface [45, 46], an adequate number of flasks was prepared to allow filtration of the entire contents of at least three flasks per sampling point. A single batch of each culture medium (control, 1% and 2% SCW) was prepared and divided into 18 × 250 *mL* Erlenmeyer flasks per condition (150 *mL* each), corresponding to a total of 54 flasks per independent experimental run. Three independent flasks per timepoint, including T0, were used as biological replicates. At T0, immediately after inoculation, the cultures had an initial biomass concentration of approximately 0.03 *g L*^−1^.

Growth was monitored by gravimetric determination of biomass dry weight. At each sampling point, the entire contents of each flask were vacuum-filtered (VacuuBrand MZ 2C, series 20547725) through pre-weighed 0.45 *μm* cellulose nitrate membrane filters (Prat Dumas, REF M0475545). Filters were dried overnight in a ventilated oven at 50°*C* and subsequently weighed using an analytical balance (Kern ABJ series, Kern & Sohn GmbH, Germany; readability 0.0001 *g*). Biomass dry weight was calculated as the difference between the final and initial filter mass and expressed as *g L*^−1^. To strengthen dataset reliability and account for experimental variability, each assay was independently repeated twice.

Biomass data are reported as mean ± standard deviation (n = 6, three flasks × two independent assay repeats), and replicate variability was assessed using the coefficient of variation (CV%), defined as the ratio between the standard deviation and the mean expressed as a percentage.

### 2.3. Mathematical model

To quantitatively describe the combined effects of light regime and SCW supplementation on biomass accumulation, a mechanistic kinetic model was developed based on three simplifying assumptions: (i) the biomass concentration at stationarity is determined by nutrient availability, independently of photoperiod; (ii) primary substrate assimilation is strictly coupled to active photosynthesis; and (iii) photoperiod modifies the relative contributions of light-dependent and whey-associated growth processes without altering the imposed stoichiometric asymptote. A fourth hypothesis— a phenomenological net dark-phase growth contribution proportional to SCW availability (*ε* > 0)—was tested as an extension to explain faster-than-predicted early growth under alternating light with SCW supplementation.

The temporal evolution of biomass *M* [*g L*^−1^], primary inorganic substrate *S* [*g L*^−1^], and secondary organic

substrate *P* [*g L*^−1^] (effective concentration of assimilable organic compounds supplied by SCW within the growth medium) is described by the following system of coupled ordinary differential equations:

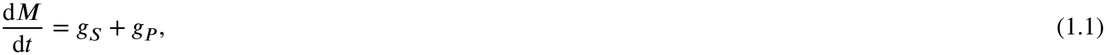

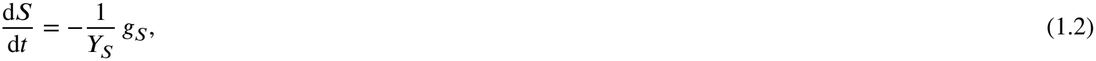

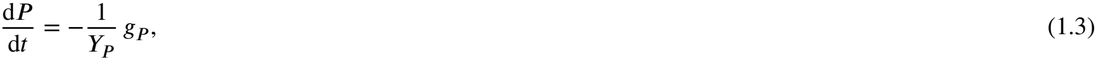

where *Y*_*S*_ and *Y*_*P*_ are the stoichiometric biomass yields on the inorganic and organic substrates, respectively. The terms *g*_*S*_ and *g*_*P*_ represent Monod-type volumetric growth rates, modulated by a binary light availability function *I*(*t*):

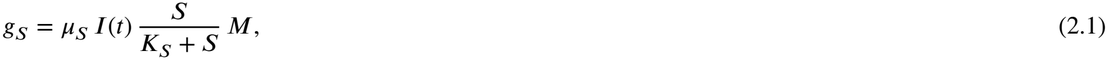

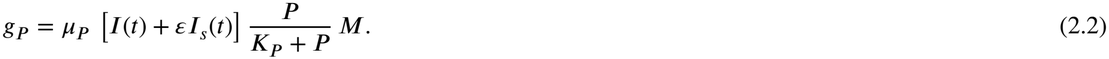

Here, *μ*_*S*_ and are maximum specific growth rates [day^−1^], *K*_*S*_ and *K*_*p*_ are half-saturation constants [*g* ^−1^], and *ε* is a phenomenological dark-phase growth multiplier. Under continuous illumination, *I*(*t*) = 1 at all times. Under alternating illumination, *I*(*t*) = 1 during the light phase (the first 16 h of each 24 h cycle) and *I*(*t*) = 0 during the dark phase. The complementary dark indicator *I*_*s*_(*t*) is defined as *I*_*s*_(*t*) = 1 − *I*(*t*) under alternating light (i.e., 1 during the 8 h dark phase and 0 otherwise), and *I*_*s*_(*t*) = 0 under continuous light.

The term *εI*_*s*_(*t*) therefore contributes only during the dark phase under alternating illumination, with a magnitude dependent on SCW availability through the Monod term *P* /(*K*_*P*_ + *P*). When *ε* = 0, the model contains no dark-phase growth contribution, whereas *ε* > 0 allows a net growth contribution during darkness. This term is phenomenological: it represents the dark-phase contribution to biomass accumulation without resolving whether it arises from direct heterotrophic assimilation of SCW organics or from consumption of glycogen accumulated photosynthetically during the light phase. Primary substrate assimilation (*g*_*S*_) remains strictly light-dependent.

The model was initialised with the measured initial biomass, *M*_0_ = 0.031 g L^−1^ (mean of the T0 measurements).

The initial inorganic substrate was set to *S*_0_ = 1.55 g L^−1^, close to the total solute mass of BG11 (Table 1). The initial SCW substrate *P*_0_ was proportional to the SCW dose: *P*_0_ = 0 in the control, 0.66 g L^−1^ at 1% v/v and 1.33 g L^−1^ at 2% v/v. These values assume that all soluble solids of SCW (6.5 °Brix, Table 2) are assimilable and that SCW has a density of 1.02 g mL^−1^ (not measured).

### 2.4. Parameterization

The stoichiometric yield coefficients *Y*_*S*_ and *Y*_*P*_ were estimated analytically from the continuous light regime: *Y*_*S*_ was derived from the unsupplemented control stationary value, and *Y*_*P*_ via linear regression of (*M*_stationary_ − *M*_control_) against SCW dose. Equivalently, *Y*_*S*_ *S*_0_ is the biomass gain of the control over the inoculum and *Y*_*P*_ *P*_0_ is the additional gain per SCW dose; these gains are obtained directly from the biomass data, and the yields follow from them by dividing by the assumed *S*_0_ and *P*_0_.

The half-saturation constants were handled as follows: *K*_*S*_ = 0.8 g L^−1^ was fixed due to identifiability limitations in the batch biomass data, while *K*_*P*_ = 0.92 g L^−1^ was determined using scalar numerical optimization (minimize_scalar). This optimization converged consistently to this value, supporting its identification as a robust minimum rather than a local artifact. Both parameters represent apparent whole-culture half-saturation constants rather than single-transporter biochemical affinities [47].

The remaining three parameters (*μ*_*S*_, *μ*_*P*_ , and *ε*) were estimated jointly via MCMC ensemble sampling implemented in Python [48]. Uniform priors were assigned to *μ*_*S*_ ∈ [0.05, 0.8] and *μ*_*P*_ ∈ [0.05, 2]. A log-normal prior was assigned to *ε* (centered at 1.7, with *σ* = 1.0 in log space, consistent with estimates from early growth ratios).

All six experimental scenarios (three SCW supplementation levels × two photoperiods) were calibrated simultaneously. Posterior distributions are summarized using the median and the 16th–84th percentile intervals, and are shown in Figure S2.

## 3. Results

### 2.1. Dose-dependent effect of SCW on biomass production

SCW supplementation increased biomass concentration at day 25 under both illumination regimes, with an approximately linear dose-dependence (Figure 1). Under continuous illumination, the observed values were close to the model-predicted stoichiometric stationary concentrations of 0.54 g L^−1^ for the control, 0.75 g L^−1^ for 1% SCW, and 0.98 g L^−1^ for 2% SCW (Figure 2). These values correspond to approximately 1.4-fold and 1.8-fold increases relative to the unsupplemented control, respectively. The dose-dependent response was described using an apparent biomass yield on SCW-associated substrate of *Y*_*P*_ = 0.33 ± 0.05 g g^−1^, compared with *Y*_*S*_ = 0.33 ± 0.03 g g^−1^ for the effective inorganic substrate.

**Figure 1.**
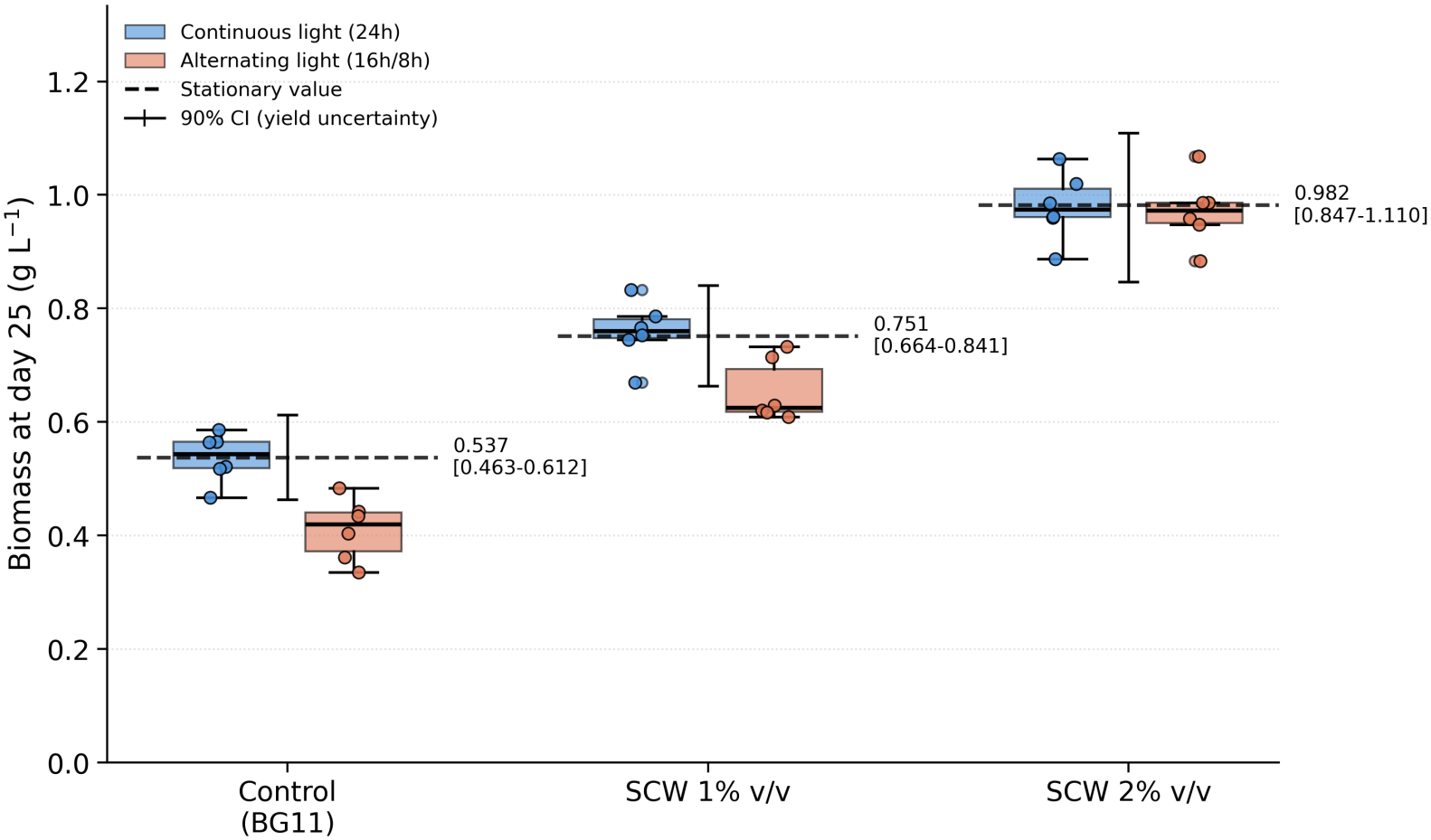
Observed biomass concentration at day 25 under continuous (blue) and alternating 16 h light/8 h dark (red) photoperiods for the control (BG11 only), 1% SCW, and 2% SCW supplementation. Boxplots show the median and interquartile range, with individual biological replicates displayed as jittered points (*n* = 6 per condition). Dashed horizontal lines indicate the model-predicted stoichiometric stationary concentrations.

**Figure 2.**
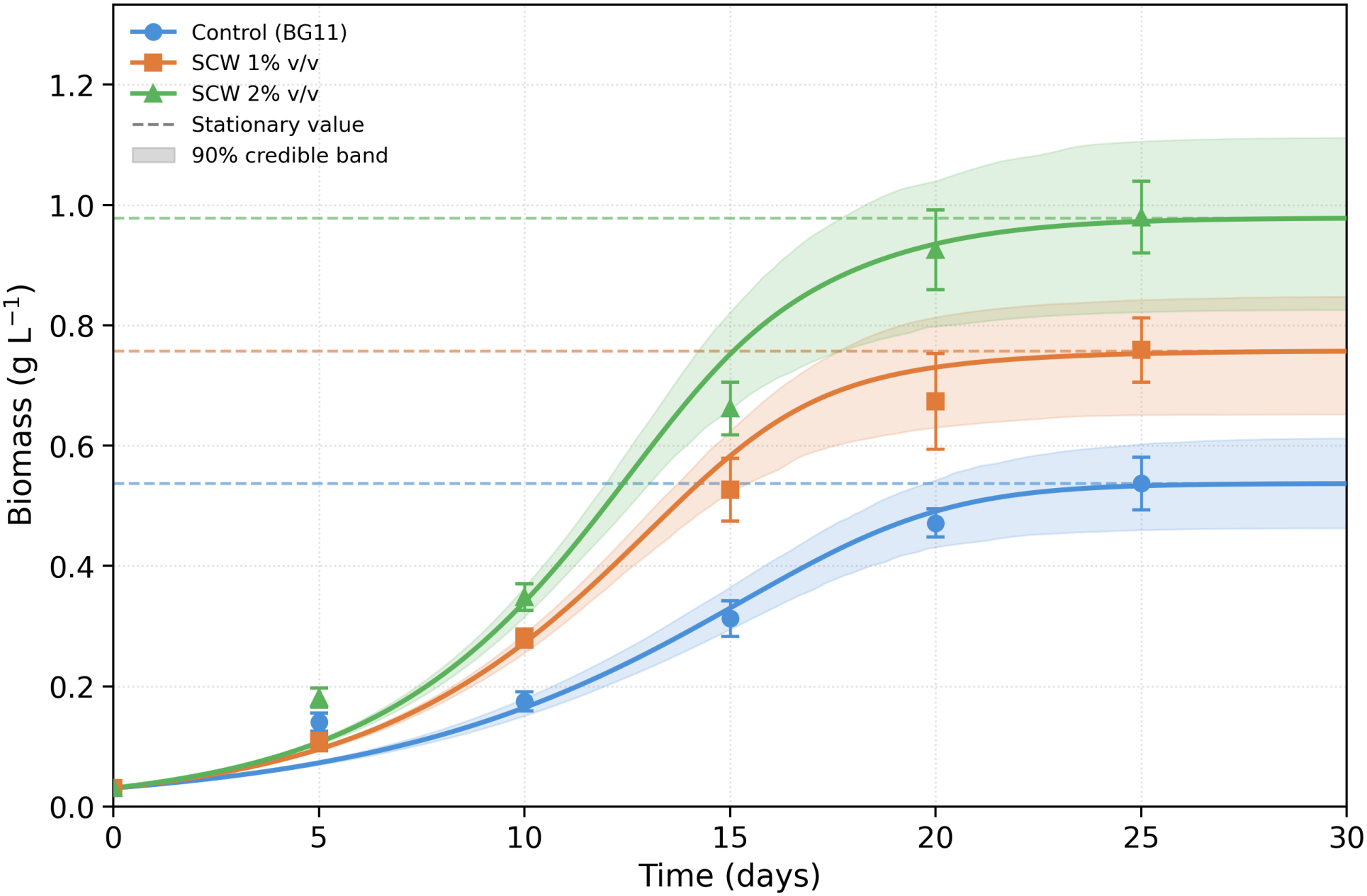
Biomass concentration of *A. platensis* under continuous illumination (24 h) for control (BG11 only, blue circles), SCW 1% v/v (orange squares), and SCW 2% v/v (green triangles). Points: observed mean ± SD (*n* = 6). Solid lines: model trajectory at posterior median parameters. Shaded bands: 90% credible interval propagating both MCMC posterior samples (*μ*_*S*_, *μ*_*P*_ , *ε*) and stoichiometric yield uncertainty (*Y*_*S*_, *Y*_*P*_). Dashed lines: model-predicted stoichiometric stationary values.

Under alternating illumination (Figure S1), the control and 1% SCW cultures remained below their model-predicted stationary concentrations at day 25 and continued to accumulate biomass during the final observation interval. In contrast, the 2% SCW treatment approached its predicted stationary concentration, with day-25 biomass comparable to that observed under continuous illumination (Figure 1).

### 3.2. Model dynamics and fit

MCMC inference yielded *μ*_*S*_ = 0.26 (± 0.01) day^−1^, *μ*_*P*_ = 0.13 (± 0.01) day^−1^, and *ε* = 3.7 (± 0.9). Posterior distributions for *μ*_*S*_, *μ*_*P*_ , and *ε* are shown in Figure S2. The model reproduced sigmoidal biomass growth dynamics across all six scenarios with a single shared parameter set (Figures 2 and S1). Model goodness of fit was assessed using the root mean square error (RMSE) computed on group means, yielding an RMSE of 0.06 *g L*^−1^ across all six calibration scenarios.

Figure 1 compares the observed and model-predicted biomass at day 25 for all six scenarios. Under alternating illumination, the control and 1% SCW cultures remained visibly below their model-predicted stoichiometric stationary values, reflecting the kinetic delay associated with the dark phase. At 1% SCW, this difference indicates that the culture had not yet reached stationary phase by day 25. In contrast, at 2% SCW, both photoperiod conditions converged closely to the same model-predicted stationary value (0.98 *g L*^−1^).

## 4. Discussion

The first key finding is that Parmigiano Reggiano SCW supplementation raises the achievable biomass ceiling of *A. platensis* in a dose-dependent, stoichiometrically predictable manner. Under continuous light, the unsupplemented control accumulated 0.54 ± 0.04 g L^−1^ of biomass above the inoculum, whereas each 1% v/v of SCW added 0.21 ± 0.03 g L^−1^ to the stationary biomass, about 40% of the control gain. This gain is consistent with mixotrophic enhancement reported for *A. platensis* on dairy substrates [49, 29], and quantitatively explained by our kinetic model: each SCW dose contributes a fixed stoichiometric increment to the biomass ceiling, both under continuous and alternating light. Under continuous light, model trajectories tracked observed growth closely within the 90% credible band for all three treatments. Under alternating light (Figure S1), the *ε* term improved early-phase fit substantially compared to the strict photoautotrophic hypothesis, consistent with a net dark-phase growth contribution that scales with SCW dose [50]. The dose-response is thus mechanistically supported, rather than simply empirical, and the model provides a quantitative tool for predicting achievable biomass concentrations under different SCW supplementation levels.

Within the tested range, the increment in the biomass ceiling was consistent with a linear dependence on SCW dose, which the model expresses through a constant apparent yield *Y*_*P*_ . With only two supplementation levels, however, this linearity is a modelling assumption rather than a tested property, and it is unlikely to hold indefinitely. At higher doses, increased turbidity and light attenuation, accumulation of inhibitory compounds, competition from heterotrophic bacteria, or limitation by a nutrient supplied by BG11 rather than by SCW may reduce the marginal gain per unit of whey. For industrial-scale applications, extending the dose range to identify where the increment departs from linearity would be particularly valuable: it would define the supplementation level that maximises biomass per unit of SCW, and the operating window within which the present model can be used for prediction.

The second key finding is that at 2% SCW, the different biomass accumulation dynamics associated with alternating 16 h/8 h illumination is largely overcome within 25 days, enabling final biomass comparable to that obtained under continuous illumination. In the model, this behaviour is captured by the dark-phase parameter *ε*, which acts as an effective multiplier of the SCW-associated growth term during darkness. Because *ε* is greater than one, it should not be interpreted as a literal fractional uptake efficiency or as a directly measured biochemical rate. Rather, it is a phenomenological parameter that summarizes the biomass gains under alternating light that cannot be explained by light-phase growth alone. This contribution remains dose-dependent through the Monod saturation term for SCW-derived organics, consistent with the faster early growth observed at the higher SCW dose under alternating light.

The distinction between kinetic limitation and physiological potential is particularly relevant when interpreting the effect of photoperiod. In photosynthetic microorganisms, the light–dark cycle should not be interpreted as a simple alternation between a productive and an inactive phase, but rather as an integrated metabolic rhythm supporting cellular homeostasis, photosynthetic regulation, and maintenance of cellular performance [51, 52]. In control cultures, biomass accumulation remained largely dependent on light-driven carbon fixation, resulting in a stronger modulation of growth kinetics under alternating illumination. Conversely, high SCW-enriched cultures achieved comparable biomass yields under alternating light/dark cycles and continuous light, suggesting an enhanced light-to-biomass conversion efficiency or a different metabolic mechanism [53]. Variation in light availability therefore primarily affects the rate of biomass accumulation rather than necessarily the biomass-forming potential of the culture. In this context, SCW supplementation may enhance the trophic flexibility of *A. platensis*, allowing the integration of organic carbon metabolism with photosynthetic activity when light availability decreases [54].

The mechanistic basis of *ε* cannot be resolved from biomass data alone. It may reflect direct heterotrophic assimilation of SCW organics during darkness, consumption of glycogen reserves accumulated photosynthetically during the light phase, or a combination of both processes. These mechanisms would produce similar signatures in biomass trajectories. Both are plausible for mixotrophic cyanobacteria [50, 20]. Measurements of substrate depletion, intracellular storage compounds, and biomass change across light–dark cycles would be required to distinguish between these alternatives.

At lower SCW doses, or without supplementation where *ε* has no effect because *P* = 0, the organic contribution is insufficient to compensate fully for the dark-phase kinetic penalty within 25 days. Because a single *Y*_*P*_ is used for both photoperiods in this model, the stoichiometric stationary value is identical for both by construction. Whether this equivalence would hold if *Y*_*P*_ were estimated independently under alternating light cannot be confirmed with the present data, since only the 2% SCW alternating-light cultures reached the stationary phase within the experimental window.

An unexpected observation at day 5 is that alternating light cultures supplemented with SCW showed higher biomass than the corresponding continuous light cultures, a pattern that reverses at later timepoints. This transient difference, which falls within replicate variability (CV 12–14%), is not yet explained by the current model and would need dedicated early-timepoint sampling (day 1–3) to test candidate explanations. Future work including substrate concentration measurements would substantially improve parameter identifiability and model transferability.

The main structural limitation of the model is that the photoperiod-independent stationary value is imposed through a single stoichiometric yield for each substrate class. This assumption is biologically plausible if photoperiod modifies the rate of substrate conversion without substantially changing the final conversion yield, but it cannot be independently verified for all alternating-light treatments in the present dataset because not all trajectories reached stationary phase within 25 days. The strongest direct support comes from the 2% SCW treatment, where continuous and alternating light converge to similar final biomass within the experimental window. For the control and 1% SCW treatments, the model should therefore be interpreted as a parsimonious kinetic explanation rather than definitive proof of an unchanged asymptote.

Beyond biomass accumulation, the increased contribution of mixotrophic metabolism raises important questions regarding cellular carbon allocation and biomass quality. Increased availability of organic carbon may influence the balance between protein synthesis, pigment maintenance, and storage compound accumulation, as reported for mixotrophic cultivation of microalgae and cyanobacteria [55]. Because biochemical composition was not evaluated in the present study, future investigations should determine whether SCW supplementation modifies the nutritional and functional properties of *A. platensis* biomass, including protein content, photosynthetic pigments, and carbon storage compounds.

From an applied perspective, the use of Parmigiano Reggiano SCW as a mixotrophic substrate aligns with circular bioeconomy principles. The finding that 2% v/v SCW largely reduces the difference in biomass accumulation dynamics between continuous and alternating illumination within the experimental window supports the potential development of cultivation systems based on natural light cycles or reduced artificial illumination schedules. However, dedicated energy and economic assessments are required to determine whether this approach can effectively reduce operating costs at the industrial scale.

## 5. Conclusions

This study demonstrates that second cheese whey (SCW), a by-product of the dairy industry, can act as an effective secondary source for *Arthrospira platensis* cultivation, increasing the achievable biomass ceiling in a dose-dependent and quantitatively predictable manner. By integrating experimental observations with a dual-substrate, light-dependent kinetic model, we show that SCW supplementation does not simply enhance biomass production, but can partially reduce the dependence of biomass accumulation on continuous light availability by providing an additional trophic pathway. The results indicate that, at 25°C, photoperiod primarily influenced the rate of biomass accumulation. Specifically, cultures supplemented with 2% v/v SCW under alternating light reached a day-25 biomass concentration comparable to that observed under continuous illumination. This finding suggests that, at the highest SCW supplementation level tested, SCW may support biomass accumulation under reduced daily light exposure. However, as the control and 1% SCW cultures under alternating light appeared not to have fully reached stationary phase within the experimental period, particularly the control, longer cultivation would be required to establish whether comparable final biomass-forming capacities can also be achieved across photoperiods under these conditions.

## Acknowledgements

L.M. thanks Lorenzo Tinacci and Jacopo Lupi for enlightning discussions.

## Fundings

This research study was funded by PR-FESR Emilia-Romagna 2021-2027, Asse 1, Azione 1.1.2. Progetto: ALGENFOR [CUP E87G22000630003]. https://www.algenfor.eu/

## CRediT authorship contribution statement

**Claudio Brandoli:** Conceptualization, Methodology, Investigation, Formal analysis, Writing – original draft, Writing – review & editing, Supervision. **Antonio Spanu:** Methodology, Formal analysis, Software, Writing – original draft, Writing – review & editing, Visualization. **Giulia Neri:** Formal analysis. **Francesca Masino:** Writing – review & editing, Funding acquisition, Project administration. **Elisabetta Sgarbi:** Methodology, Writing – review & editing. **Leonardo Miele:** Conceptualization, Methodology, Formal analysis, Writing – original draft, Writing – review & editing, Supervision.

## Notes

### Competing Interest Statement

The authors have declared no competing interest.

## References

[1] M. D. Guiry, G. M. Guiry AlgaeBase, World-wide electronic publication, National University of Ireland, Galway, 2024. URL: https://www.algaebase.org, database record dated 15 May 2024; available at https://www.algaebase.org; accessed 16 September 2026.

[2] C. H. G. Jung, S. Braune, P. Waldeck, J.-H. Küpper, I. Petrick, F. Jung, Morphology and growth of Arthrospira platensis during cultivation in a flat-type bioreactor, Life 11 (2021) 536.

[3] L. S. Ferreira, M. S. Rodrigues, A. Converti, S. Sato, J. C. M. Carvalho, Kinetic and growth parameters of Arthrospira (Spirulina) platensis cultivated in tubular photobioreactor under different cell circulation systems, Biotechnology and Bioengineering 109 (2012) 444–450.

[4] M. C. Matsudo, R. P. Bezerra, S. Sato, A. Converti, J. C. M. de Carvalho, Photosynthetic efficiency and rate of CO2 assimilation by Arthrospira (Spirulina) platensis continuously cultivated in a tubular photobioreactor, Biotechnology Journal 7 (2012) 1412–1417.

[5] A. Vonshak, Spirulina platensis (Arthrospira): Physiology, Cell-Biology and Biotechnology, CRC Press, 1997.

[6] L. J. Stal, Cyanobacteria: diversity and versatility, clues to life in extreme environments, in: Algae and Cyanobacteria in Extreme Environments, Springer, 2007, pp. 659–680.

[7] M. M. A. Mahmoud, M. M. M. El-Lamie, O. E. Kilany, A. A. Dessouki, Spirulina (Arthrospira platensis) supplementation improves growth performance, feed utilization, immune response, and relieves oxidative stress in nile tilapia (Oreochromis niloticus) challenged with Pseudomonas fluorescens, Fish & Shellfish Immunology 72 (2018) 291–300.

[8] A. L. Lupatini, L. M. Colla, C. Canan, E. Colla, Potential application of microalga Spirulina platensis as a protein source, Journal of the Science of Food and Agriculture 97 (2017) 724–732.

[9] A. Lestingi, M. Alagawany, A. Di Cerbo, G. Crescenzo, C. Zizzadoro, Spirulina (Arthrospira platensis) used as functional feed supplement or alternative protein source: A review of the effects of different dietary inclusion levels on production performance, health status, and meat quality of broiler chickens, Life 14 (2024) 1537.

[10] S. K. Ali, A. M. Saleh, Spirulina—an overview, International Journal of Pharmacy and Pharmaceutical Sciences 4 (2012) 9–15.

[11] N. T. Eriksen, Production of phycocyanin—a pigment with applications in biology, biotechnology, foods and medicine, Applied Microbiology and Biotechnology 80 (2008) 1–14.

[12] M. Raeisossadati, O. Bumandalai, N. R. Moheimani, Can anaerobically digested food effluent support Arthrospira platensis cultivation in open ponds?, Applied Sciences 15 (2025) 3115.

[13] K. Soma, J. Kals, M. A. Opiyo, A. Ndambi, R. García-Cubero, M. J. Barbosa, E. Rurangwa, A. Vernooij, Toward sustainable food systems: can spirulina (Arthrospira platensis) become a sustainable source of protein to enhance the nutritional benefits of cultured nile tilapia (Oreochromis niloticus)?, Frontiers in Sustainable Food Systems 8 (2024) 1283150.

[14] S. Braune, C. G. Jung, J.-H. Küpper, F. Jung, Arthrospira platensis as protein-rich source for human nutrition, Life 15 (2025) 1789.

[15] V. Vannini, W. M. J. Achten, Comparative life cycle assessment of industrial and artisanal spirulina production systems, Science of The Total Environment 997 (2025) 180184.

[16] R. A. Soni, K. Sudhakar, R. S. Rana, Spirulina—from growth to nutritional product: A review, Trends in Food Science & Technology 69 (2017) 157–171.

[17] N. Sumprasit, N. Wagle, N. Glanpracha, A. P. Annachhatre, Biodiesel and biogas recovery from Spirulina platensis, International Biodeterioration & Biodegradation 119 (2017) 196–204.

[18] F. Arahou, I. Lijassi, A. Wahby, L. Rhazi, M. Arahou, I. Wahby, Spirulina-based biostimulants for sustainable agriculture: Yield improvement and market trends, BioEnergy Research 16 (2023) 1401–1416.

[19] S. d. P. Ramos, M. Bürck, S. F. F. da Costa, M. Assis, A. R. C. Braga, Spirulina as a key ingredient in the evolution of eco-friendly cosmetics, BioTech 14 (2025) 41.

[20] K. Chojnacka, A. Noworyta, Evaluation of Spirulina sp. growth in photoautotrophic, heterotrophic and mixotrophic cultures, Enzyme and Microbial Technology 34 (2004) 461–465.

[21] P. Mishra, S. M. Prasad, Low dose UV-B radiation induced mild oxidative stress impact on physiological and nutritional competence of Spirulina (Arthrospira) species, Plant Stress 2 (2021) 100039.

[22] W. Blanken, M. Cuaresma, R. H. Wijffels, M. Janssen, Cultivation of microalgae on artificial light comes at a cost, Algal Research 2 (2013) 333–340.

[23] J. Zhai, X. Li, W. Li, M. H. Rahaman, Y. Zhao, B. Wei, H. Wei, Optimization of biomass production and nutrients removal by Spirulina platensis from municipal wastewater, Ecological Engineering 108 (2017) 83–92.

[24] M. Bürck, C. Fratelli, M. C. A. de Amarante, A. R. C. Braga, Unveiling the potential of spirulina biomass—a glimpse into the future circular economy using green and blue ingredients, Biomass 4 (2024) 704–719.

[25] B. Barati, Y. Li, S. Gusev, D. P. L. Rousseau, S. W. H. Van Hulle, Unlocking the potential of brewery wastewater: Sustainable cultivation of Arthrospira platensis for biomass and phycocyanin production, Journal of Water Process Engineering 70 (2025) 107107.

[26] G. W. Smithers, Whey and whey proteins—from gutter-to-gold, International Dairy Journal 18 (2008) 695–704.

[27] F. Carvalho, A. R. Prazeres, J. Rivas, Cheese whey wastewater: Characterization and treatment, Science of the Total Environment 445–446 (2013) 385–396.

[28] V. Athanasiadou, E. E. Klontza, P. Dimitriou-Christidis, M. Fountoulakis, D. F. Lekkas, Evaluation of Arthrospira (Spirulina) platensis growth on cheese whey in the context of circular economy, Sustainable Chemistry and Pharmacy 34 (2023) 101173.

[29] A. Cavallini, S. Torre, L. Usai, M. Casula, G. Fais, P. Nieri, A. Concas, G. A. Lutzu, Effect of cheese whey on phycobiliproteins production and FAME profile by Arthrospira platensis (Spirulina): Promoting the concept of a circular bio-economy, Sustainable Chemistry and Pharmacy 40 (2024) 101625.

[30] E. Sventzouri, K. Pispas, G. G. Kournoutou, M. Geroulia, E. Giakoumatou, S. S. Ali, M. Kornaros, Evaluation of growth performance, biochemical composition, and polyhydroxyalkanoates production of four cyanobacterial species grown in cheese whey, Microorganisms 13 (2025) 1157.

[31] A. F. Pires, N. G. Marnotes, O. D. Rubio, A. C. Garcia, C. D. Pereira, Dairy by-products: A review on the valorization of whey and second cheese whey, Foods 10 (2021) 1067.

[32] ISMEA, Parmigiano Reggiano –scheda prodotto DOP-IGP, ISMEA Mercati, online product data sheet, 2025. Available at https://www.ismeamercati.it/flex/FixedPages/IT/FoodCertificatoScheda.php/L/IT/ID/146/REG/8/tip/2/from/-; accessed 1 July 2026.

[33] Regione Emilia-Romagna, Circular economy, Regione Emilia-Romagna, Transizione ecologica portal, 2026. Available at https://imprese.regione.emilia-romagna.it/transizione-ecologica/normativa-strategia-pianificazione/strategia/circular-economy; accessed 1 July 2026.

[34] O. Bernard, Hurdles and challenges for modelling and control of microalgae for CO2 mitigation and biofuel production, Journal of Process Control 21 (2011) 1378–1389.

[35] G. Cogne, J.-B. Gros, C.-G. Dussap, Identification of a metabolic network structure representative of Arthrospira (Spirulina) platensis metabolism, Biotechnology and Bioengineering 84 (2003) 667–676.

[36] N. Aowtrakool, A. Sopitthummakhun, T. Laomettachit, M. Ruengjitchatchawalya, A dynamic model for predicting biomass and phycocyanin yields in Arthrospira (Spirulina) platensis: A guidance for effective batch cultivation, Algal Research 83 (2024) 103709.

[37] H. Y. Lee, L. E. Erickson, S. S. Yang, Kinetics and bioenergetics of light-limited photoautotrophic growth of Spirulina platensis, Biotechnology and Bioengineering 29 (1987) 832–843.

[38] D. A. Marrez, M. M. Naguib, Y. Y. Sultan, Z. Y. Daw, A. M. Higazy, Evaluation of chemical composition for Spirulina platensis in different culture media, Research Journal of Pharmaceutical, Biological and Chemical Sciences 5 (2014) 1161–1171.

[39] Y. M. A. Fagiri, A. Salleh, S. A. F. El-Nagerabi, Impact of physico-chemical parameters on the physiological growth of Arthrospira (Spirulina platensis) exogenous strain UTEXLB2340, African Journal of Biotechnology 12 (2013) 5458–5465.

[40] Z. Li, Y. Liu, T. Zhou, L. Cao, Y. Cai, Y. Wang, X. Cui, H. Yan, R. Ruan, Q. Zhang, Effects of culture conditions on the performance of Arthrospira platensis and its production of exopolysaccharides, Foods 11 (2022) 2020.

[41] R. P. Bezerra, E. Y. O. Montoya, S. Sato, P. Perego, J. C. M. de Carvalho, A. Converti, Effects of light intensity and dilution rate on the semicontinuous cultivation of Arthrospira (Spirulina) platensis: A kinetic Monod-type approach, Bioresource Technology 102 (2011) 3215–3219.

[42] L. H. Pelizer, E. D. G. Danesi, C. d. O. Rangel, C. E. N. Sassano, J. C. M. de Carvalho, S. Sato, I. O. Moraes, Influence of inoculum age and concentration in Spirulina platensis cultivation, Journal of Food Engineering 56 (2003) 371–375.

[43] J. Folch, M. Lees, G. H. Sloane Stanley, A simple method for the isolation and purification of total lipides from animal tissues, Journal of Biological Chemistry 226 (1957) 497–509.

[44] O. H. Lowry, N. J. Rosebrough, A. L. Farr, R. J. Randall, Protein measurement with the Folin phenol reagent, Journal of Biological Chemistry 193 (1951) 265–275.

[45] M. B. F. Silva, E. G. Azero, C. M. L. L. Teixeira, C. T. Andrade, Influence of culture conditions on the production of extracellular polymeric substances (EPS) by Arthrospira platensis, Bioresources and Bioprocessing 7 (2020) 47.

[46] X. Wang, X. Wen, Y. Li, Y. Zhang, Y. Zhao, Z. Tang, D. Yu, J. Li, B. Ge, The response of extracellular polymeric substances (EPS) in Spirulina platensis to environmental stresses: salt, light and heavy metal, Environmental Research 285 (2025) 122513.

[47] S.-i. Maeda, T. Omata, Substrate-binding lipoprotein of the cyanobacterium Synechococcus sp. strain PCC 7942 involved in the transport of nitrate and nitrite, Journal of Biological Chemistry 272 (1997) 3036–3041.

[48] D. Foreman-Mackey, D. W. Hogg, D. Lang, J. Goodman, emcee: The MCMC hammer, Publications of the Astronomical Society of the Pacific 125 (2013) 306–312.

[49] M. I. B. Pereira, B. M. E. Chagas, R. Sassi, G. F. Medeiros, E. M. Aguiar, L. H. F. Borba, E. P. E. Silva, J. C. Andrade Neto, A. H. N. Rangel, Mixotrophic cultivation of Spirulina platensis in dairy wastewater: Effects on the production of biomass, biochemical composition and antioxidant capacity, PLOS ONE 14 (2019) e0224294.

[50] A. Vonshak, S. M. Cheung, F. Chen, Mixotrophic growth modifies the response of Spirulina (Arthrospira) platensis (Cyanobacteria) cells to light, Journal of Phycology 36 (2000) 675–679.

[51] G. Pattanayak, M. J. Rust, The cyanobacterial clock and metabolism, Current Opinion in Microbiology 18 (2014) 90–95.

[52] D. G. Welkie, B. E. Rubin, S. Diamond, R. D. Hood, D. F. Savage, S. S. Golden, A hard day’s night: Cyanobacteria in diel cycles, Trends in Microbiology 27 (2019) 231–242.

[53] M. A. Furmaniak, A. E. Misztak, M. D. Franczuk, A. Wilmotte, M. Waleron, K. F. Waleron, Edible cyanobacterial genus Arthrospira: Actual state of the art in cultivation methods, genetics, and application in medicine, Frontiers in Microbiology 8 (2017) 2541.

[54] S. Lucius, M. Hagemann, The primary carbon metabolism in cyanobacteria and its regulation, Frontiers in Plant Science 15 (2024) 1417680.

[55] T. Castillo, D. Ramos, T. García-Beltrán, M. Brito-Bazan, E. Galindo, Mixotrophic cultivation of microalgae: An alternative to produce high-value metabolites,Biochemical Engineering Journal 176 (2021) 108183.

