## Supplemental Figures for "A midnight snack for spirulina: a dual-substrate kinetic model for second cheese whey supplementation of *Arthrospira platensis*"

### Supplementary Material

#### A midnight snack for spirulina: a dual-substrate kinetic model reveals how second cheese whey shapes biomass production in *Arthrospira platensis*

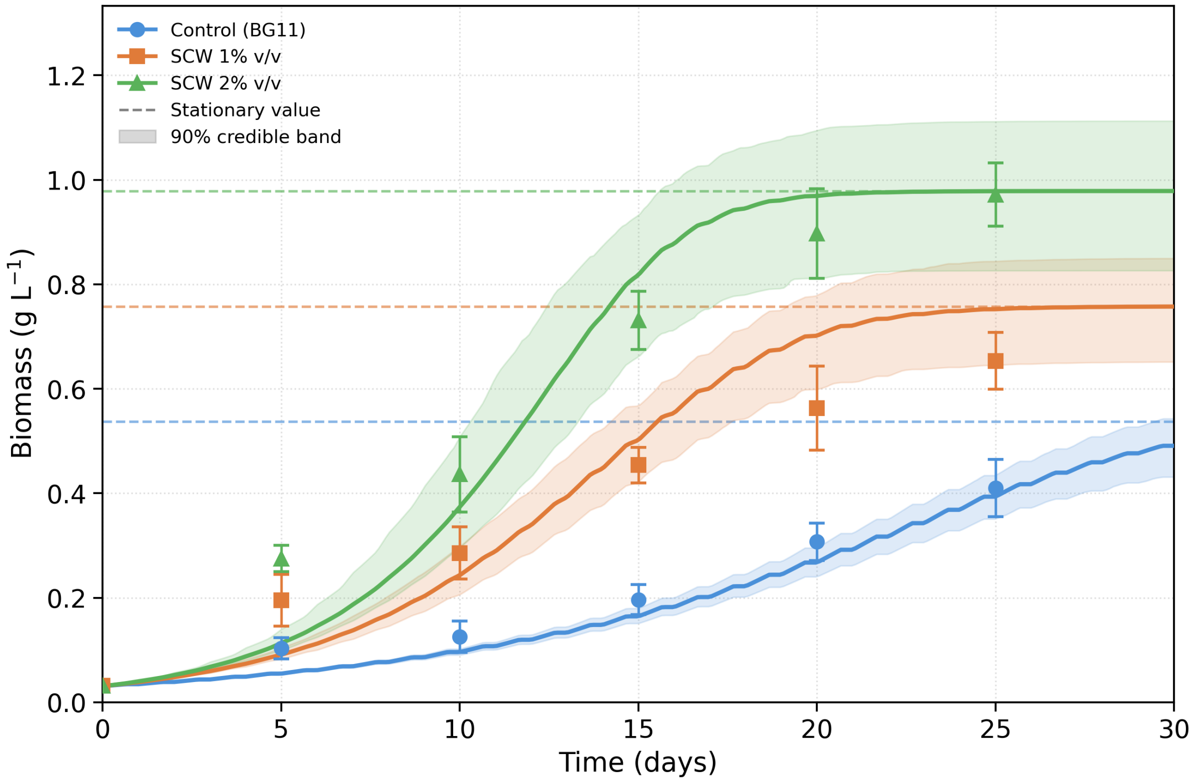


Figure S1: Biomass concentration of *A. platensis* under alternating illumination (16 h light/8 h dark) for the control (BG11 only), SCW 1% v/v, and SCW 2% v/v treatments. Points represent observed mean ± SD ($n = 6$). Solid lines indicate model trajectories at posterior median parameters, shaded areas represent 90% credible intervals, and dashed horizontal lines indicate model-predicted stoichiometric stationary values. The kinetic delay associated with alternating illumination is visible at day 25 for the control and SCW 1% treatments.

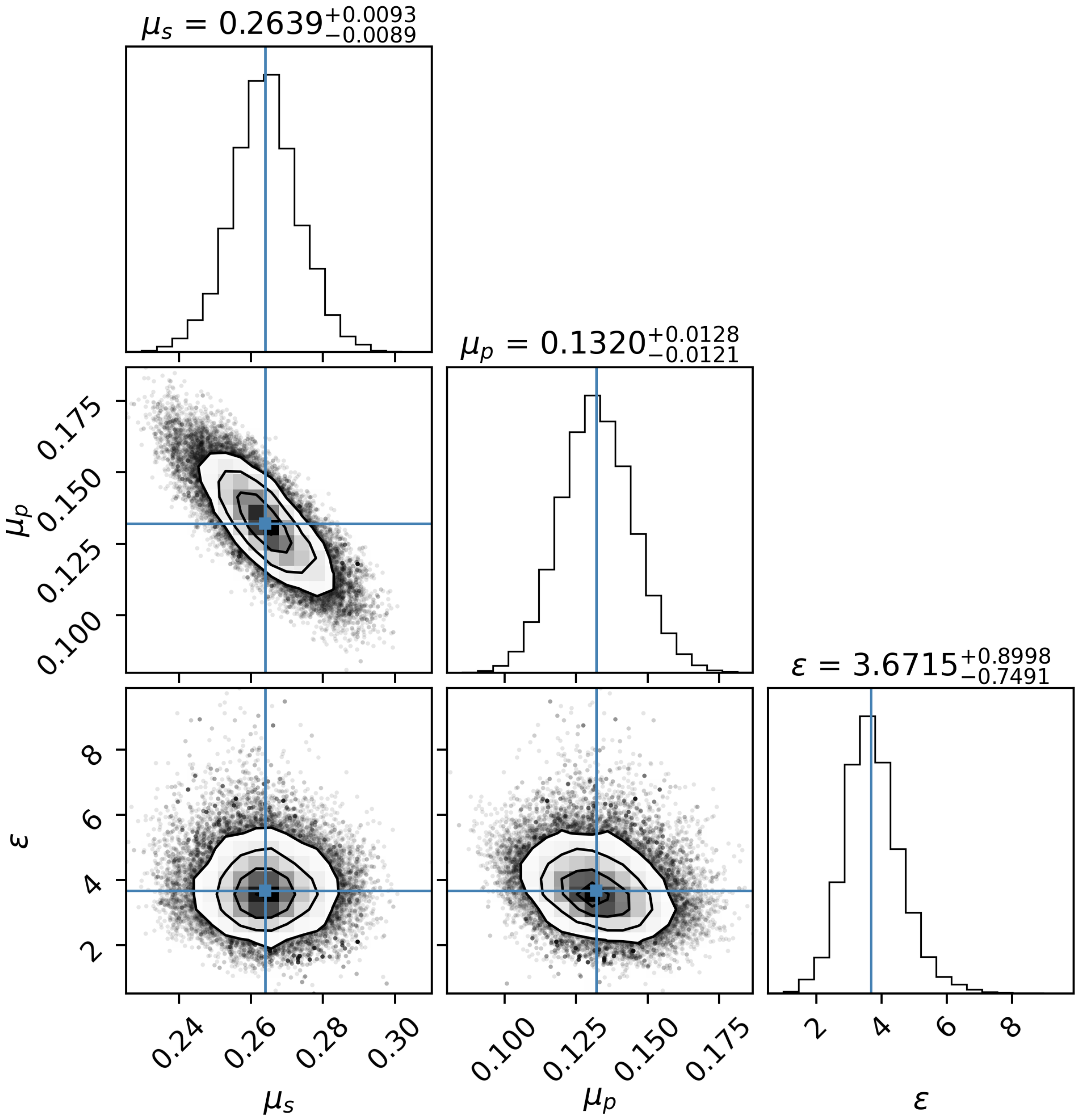


Figure S2: Posterior distributions and pairwise correlations of the kinetic parameters $\mu_S$ and $\mu_P$ and the phenomenological dark-phase multiplier $\varepsilon$, estimated by joint Markov chain Monte Carlo (MCMC) inference. Blue lines indicate posterior medians.
